# pyaging: a Python-based compendium of GPU-optimized aging clocks

**DOI:** 10.1101/2023.11.28.569069

**Authors:** Lucas Paulo de Lima Camillo

## Abstract

**Motivation:** Aging is intricately linked to diseases and mortality and is reflected in molecular changes across various tissues. The development and refinement of biomarkers of aging, healthspan, and lifespan using machine learning models, known as aging clocks, leverage epigenetic and other molecular signatures. Despite advancements, as noted by the Biomarkers of Aging Consortium, the field grapples with challenges, notably the lack of robust software tools for integrating and comparing these diverse models.

**Results:** I introduce `pyaging`, a comprehensive Python package, designed to bridge the gap in aging research software tools. `pyaging` integrates over 30 aging clocks, with plans to expand to more than 100, covering a range of molecular data types including DNA methylation, transcriptomics, histone mark ChIP-Seq, and ATAC-Seq. The package features a variety of model types, from linear and principal component models to neural networks and automatic relevance determination models. Utilizing a PyTorch-based backend for GPU acceleration, `pyaging` ensures rapid inference even with large datasets and complex models. The package supports multi-species analysis, currently including humans, various mammals, and C. elegans.

**Availability and Implementation:** `pyaging` is accessible at https://github.com/rsinghlab/pyaging. The package is structured to facilitate ease of use and integration into existing research workflows, supporting the flexible anndata data format.

**Supplementary Information:** Supplementary materials, including detailed documentation and usage examples, are available online at the `pyaging` documentation site (https://pyaging.readthedocs.io/en/latest/index.html).

## Introduction

As we entered the 21st century, longevity studies became the cornerstone of aging research in various model organisms. The span of these studies ranged from a few days in Caenorhabditis elegans to several weeks in Drosophila melanogaster, extending up to a few years in Mus musculus. This spectrum allowed for manageable daily mortality tracking in the burgeoning field of gerontology. Nonetheless, the feasibility of lifespan studies, both in terms of time and cost, remained a significant challenge. The transformative work by Horvath in 2013 [1] marked a pivotal moment, introducing a reliable age predictor and catalyzing a new domain of research focused on the development and refinement of biomarkers of aging, healthspan, and lifespan.

Presently, the field boasts over a hundred aging clocks—machine learning models designed to predict various aspects of aging. DNA methylation, undoubtedly the most popular data type for constructing aging clocks, is complemented by other molecular signatures like transcriptomics, proteomics, blood chemistry, histone modification, and chromatin accessibility, each offering unique advantages. However, there exists a notable gap in software tools that consolidate these diverse aging clocks for comparative analysis. A few notable initiatives, such as the R packages methylclock and methylcypher [2, 3], represent steps towards addressing this need.

Yet, the development of aging biomarkers is not without its challenges, as underscored in a recent perspective [4]. Current software tools in this domain face several limitations: (1) the prevalent use of R, while popular in biology, often lacks the versatility needed for complex models like neural networks, which are more effectively implemented in languages such as Python; (2) most existing age prediction packages are limited to a handful of clocks, far fewer than the actual breadth of available models; (3) a focus predominantly on DNA methylation biomarkers narrows the scope for cross-comparison across different molecular layers; (4) the lack of advanced modeling techniques, such as principal component analysis [5] and neural network-based approaches like AltumAge [6]; (5) the reliance on CPU processing results in slower inference, especially with larger datasets and more complex models; (6) a species-specific focus, predominantly on Homo sapiens.

Addressing these challenges, I have developed `pyaging`, a Python-based package that acts as a comprehensive repository for various aging clocks. `pyaging` offers: (1) a Python-centric approach, utilizing the versatile anndata data format; (2) an expanding repository, currently encompassing 30 clocks with an aim to include over 100; (3) clocks based on a diverse range of data types, encompassing DNA methylation, transcriptomics [7], histone mark ChIP-Seq [8], and ATAC-Seq [9]; (4) a variety of models, including linear, principal component linear models, neural networks, and automatic relevance determination models; (5) a PyTorch-based backend that leverages GPU processing for enhanced inference speeds; (6) a multi-species scope, currently covering human, various mammalian species, and C. elegans, with plans to integrate murine-specific clocks shortly.

## Methods

The development of the `pyaging` package commenced with an extensive review of the literature to identify a diverse array of aging clocks, encompassing various data types, computational models, and species, as summarized in Table 1.

**Table 1:** Overview of aging clocks currently available on pyaging. “PC” stands for principal components, whereas “ARD” means automatic relevance determination.

| Clock name | Species | Model type | Data type | Year | Citation |
| --- | --- | --- | --- | --- | --- |
| mammalianlifespan | multi | linear | methylation | 2023 | [10] |
| panhistone | Homo sapiens | pc-ard | histone mark | 2023 | [8] |
| h3k9me3 | Homo sapiens | pc-ard | histone mark | 2023 | [8] |
| h3k9ac | Homo sapiens | pc-ard | histone mark | 2023 | [8] |
| h3k4me3 | Homo sapiens | pc-ard | histone mark | 2023 | [8] |
| h3k4me1 | Homo sapiens | pc-ard | histone mark | 2023 | [8] |
| h3k36me3 | Homo sapiens | pc-ard | histone mark | 2023 | [8] |
| h3k27me3 | Homo sapiens | pc-ard | histone mark | 2023 | [8] |
| h3k27ac | Homo sapiens | pc-ard | histone mark | 2023 | [8] |
| mammalian3 | multi | linear | methylation | 2023 | [11] |
| mammalian2 | multi | linear | methylation | 2023 | [11] |
| mammalian1 | multi | linear | methylation | 2023 | [11] |
| ocampoatac2 | Homo sapiens | linear | atac | 2023 | [9] |
| ocampoatac1 | Homo sapiens | linear | atac | 2023 | [9] |
| dunedinpace | Homo sapiens | linear | methylation | 2022 | [12] |
| pcskinandblood | Homo sapiens | pc-linear | methylation | 2022 | [5] |
| pcphenoage | Homo sapiens | pc-linear | methylation | 2022 | [5] |
| pchorvath2013 | Homo sapiens | pc-linear | methylation | 2022 | [5] |
| pchannum | Homo sapiens | pc-linear | methylation | 2022 | [5] |
| pcgrimage | Homo sapiens | pc-linear | methylation | 2022 | [5] |
| pcdnamtl | Homo sapiens | pc-linear | methylation | 2022 | [5] |
| altumage | Homo sapiens | neural network | methylation | 2022 | [6] |
| replitali | Homo sapiens | linear | methylation | 2022 | [13] |
| bitage | C elegans | linear | transcriptomics | 2021 | [7] |
| dnamtl | Homo sapiens | linear | methylation | 2019 | [14] |
| pedbe | Homo sapiens | linear | methylation | 2019 | [15] |
| skinandblood | Homo sapiens | linear | methylation | 2018 | [16] |
| phenoage | Homo sapiens | linear | methylation | 2018 | [17] |
| horvath2013 | Homo sapiens | linear | methylation | 2013 | [1] |
| hannum | Homo sapiens | linear | methylation | 2013 | [18] |

Each identified model was then adapted to a PyTorch backend to facilitate GPU-accelerated computations. This adaptation leverages the fact that most aging clocks can be fundamentally represented through a series of matrix multiplications. Consider the most prevalent model type in aging research: linear models. In algebraic terms, with a set of coefficients ***β***, bias *ϵ*, independent variables **x**, and the dependent variable *y*, the linear model is expressed as:

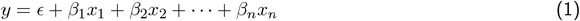

This can be elegantly reformulated using matrix notation:

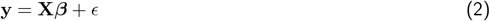

where **X** is the matrix of independent variables. Similarly, principal component-based clocks can be represented as:

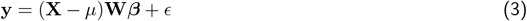

Here, *µ* denotes the centering vector for PCA, and **W** represents the rotation matrix. Thus, a broad spectrum of aging clocks are fundamentally reducible to matrix multiplications.

The implementation of age prediction in `pyaging` begins with preprocessing the data matrix. Missing values are imputed using methodologies ranging from simple mean imputation to more sophisticated techniques like KNN imputation. The input matrix is then curated to retain only the features pertinent to the selected clock, with any absent features being substituted with zeros. This approach is adopted to accommodate the diversity of data types handled by `pyaging`, and users are duly alerted of such substitutions. Additional preprocessing steps are tailored to specific models, such as scaling for AltumAge and binarization for BiTAge. The processed data is then fed into the model for age prediction. Postprocessing steps, such as anti-log-linear transformation for certain clocks like Horvath’s 2013 and the SkinAndBlood clocks, are applied as necessary. All computations are conducted using PyTorch tensors within anndata objects, ensuring efficient and scalable processing. The output includes the predicted values across all selected clocks, accompanied by the respective metadata, such as citations, for user reference.

Detailed tutorials and use-case examples are available on the documentation website: https://readthedocs.org/projects/pyaging/builds/22654195/. The code is also available on GitHub: https://github.com/rsinghlab/pyaging.

## Results

To demonstrate the capabilities of the `pyaging` package, it was applied to analyze 19 methylation aging clocks using data from AltumAge [6] and Horvath’s mammalian dataset [11]. AltumAge’s dataset comprises approximately 13,000 samples across 142 datasets, featuring beta values from Illumina’s 27k and 450k arrays. In contrast, Horvath’s mammalian dataset includes around 15,000 samples from Illumina’s custom mammalian array. My approach centered on generating correlation matrices through Spearman’s correlation, followed by examining the principal component loadings of each clock within the datasets.

Analysis of the AltumAge data, as depicted in Figure 1a, revealed that clocks measuring cell replication indicators, such as DNA telomere length and population doublings, tended to cluster. Notably, PCGrimAge showed a clustering pattern with chronological age predictors like Horvath2013 and AltumAge. Additionally, DunedinPACE [12] predominantly aligned with the mammalian predictor of relative age, mammalian2. In terms of principal component loadings, PCHannum and Hannum were the primary contributors to the variance in PC1 (Figure 1b).

**Figure 1:**
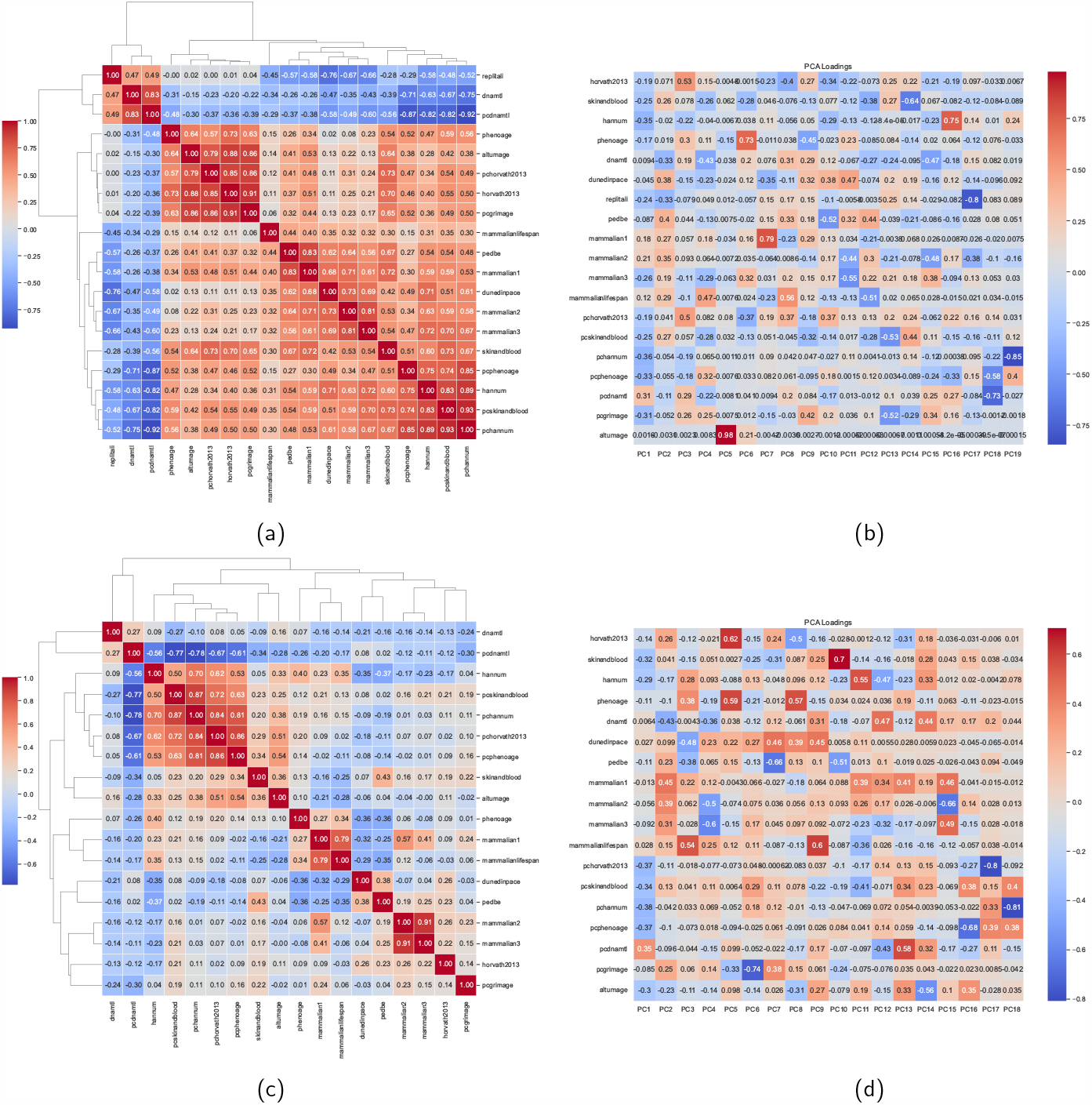
Comparisons between 19 methylation aging clocks in the AltumAge data set (a, b) and the mammalian data set (c, d). (a, c) Heatmap of Spearman’s correlation matrix for the predicted values. (b, d) Principal component loadings of the clocks explaining the variance in the predicted value matrix. of current tools, offering greater flexibility for complex models. The incorporation of a wide array of aging clocks, covering various molecular signatures, reflects the commitment to a comprehensive understanding of aging. Additionally, `pyaging` integrates advanced modeling techniques and leverages GPU processing for enhanced computational efficiency. Its multi-species capability extends its utility across a range of gerontological studies. Overall, `pyaging` not only marks a substantial progress in the field of aging biomarker research but also sets a foundation for further scientific inquiries in this rapidly developing domain. To include your own aging clock in `pyaging`, just reach out and I will aim to expand the package in a timely manner.

Conversely, the analysis of the mammalian data indicated fewer positive correlations among the clocks (Figure 1c). The most substantial cluster consisted of PC clocks, excluding PCGrimAge. It is also significant that Horvath2013 exhibited minimal correlation with other clocks in this dataset. Similar to the AltumAge data, PCHannum was the predominant factor in explaining the variance in PC1 from the mammalian dataset (Figure 1d).

A significant and obvious limitation encountered in this analysis was the substantial proportion of missing features in several clocks. For example, RepliTali was excluded from the mammalian array analysis due to a complete lack of overlapping CpG sites with the mammalian array. Similarly, principal component clocks demonstrated considerable feature deficits in the mammalian array. Despite these limitations, the exploratory analysis facilitated intriguing comparative insights. The main takeaway is the ease of use and speed of the package in handling large datasets.

## Conclusion

Despite the abundance of aging clocks developed, a critical gap remains in integrating these diverse models for comprehensive analysis, a need only partially addressed by existing tools like methylclock and methylcypher. My contribution, the `pyaging` package, represents a significant advancement in addressing these challenges. By adopting a Python-centric approach, `pyaging` overcomes the limitations inherent in the R-dominated landscape

